# Identification of genes involved in the differentiation of R7y and R7p photoreceptor cells in *Drosophila*

**DOI:** 10.1101/748095

**Authors:** James B Earl, Lauren A Vanderlinden, Laura M Saba, Steven G Britt

**Affiliations:** Department of Cell and Developmental Biology, School of Medicine, University of Colorado Anschutz Medical Campus, Aurora, CO, 80045; Department of Biostatistics and Informatics, Colorado School of Public Health, Aurora, CO 80045; Department of Pharmaceutical Sciences, Skaggs School of Pharmacy and Pharmaceutical Sciences, University of Colorado Anschutz Medical Campus, Aurora, CO 80045; Department of Neurology, Department of Ophthalmology, Dell Medical School, University of Texas at Austin, Austin, TX 78712

**Keywords:** Photoreceptor, Cell fate, Gene inactivation, Ectopic expression, Mutant screen

## Abstract

The R7 and R8 photoreceptor cells of the *Drosophila* compound eye mediate color vision. Throughout the majority of the eye, these cells occur in two principal types of ommatidia. Approximately 35% of ommatidia are of the pale type and express Rh3 in R7 cells and Rh5 in R8 cells. The remaining 65% are of the yellow type and express Rh4 in R7 cells and Rh6 in R8 cells. The specification of an R8 cell in a pale or yellow ommatidium depends on the fate of the adjacent R7 cell. However, pale and yellow R7 cells are specified by a stochastic process that requires the genes *spineless*, *tango* and *klumpfuss*. To identify additional genes involved in this process we performed a genetic screen using a collection of 480 *P{EP}* transposon insertion strains. We identified genes that when inactivated and/or ectopically expressed in R7 cells resulted in a significantly altered percentage of Rh3 expressing R7 cells (Rh3%) from wild-type. 53 strains resulted in altered Rh3% in the heterozygous inactivation arm of the screen. 36 strains resulted in altered Rh3% in the ectopic expression arm of the screen, where the P{EP} insertion strains were crossed to a *sevEP-GAL4* driver line. 4 strains showed differential effects between the two screens. Analyses of these results suggest that R7 cell fate specification is sensitive to perturbations in transcription, growth inhibition, glycoprotein ligand binding, WNT signaling, ubiquitin protease activity and Ser/Thr kinase activity, among other diverse signaling and cell biological processes.

## Introduction

Color vision in most organisms is dependent upon the expression of spectrally distinct visual pigments (opsins) in different photoreceptor cells (Jacobs 1981; Nathans *et al.* 1986; Wikler and Rakic 1990). The organization of the retinal mosaic reflects a variety of different developmental mechanisms, including regional specialization, stochastic, and precise cell-cell adjacency (Viets *et al.* 2016). *Drosophila melanogaster* is capable of color vision and is a useful experimental system for examining the developmental programs that produce photoreceptor cells having different color sensitivities (Quinn *et al.* 1974; Spatz *et al.* 1974; Chou *et al.* 1996; Chou *et al.* 1999; Tang and Guo 2001; Cook *et al.* 2003; Wernet *et al.* 2003; Mikeladze-Dvali *et al.* 2005). The compound eye consists of ∼800 ommatidia, which each contain a pair of R7 and R8 photoreceptors cells that mediate polarization sensitivity and color vision (Fortini and Rubin 1991; Yamaguchi *et al.* 2010).

Two main ommatidial subtypes were identified based on pale or yellow fluorescence when illuminated with blue light (Kirschfeld *et al.* 1978; Franceschini *et al.* 1981), and contain R7pale/R8pale (R7p/R8p) or R7yellow/R8yellow (R7y/R8y) cell pairs expressing Rh3/Rh5 or Rh4/Rh6, respectively (Chou *et al.* 1996; Papatsenko *et al.* 1997; Chou *et al.* 1999). The R7y and R7p photoreceptor cells are distributed randomly (Bell *et al.* 2007) and thought to be generated stochastically through the action of *spineless (ss)*, *tango* (*tgo*) and *klumpfuss (klu)* (Wernet *et al.* 2006; Thanawala *et al.* 2013; Johnston and Desplan 2014; Anderson *et al.* 2017). The hypothesis behind this stochastic cell-fate mechanism is that variation in the expression of *ss* in individual R7 cells in the developing pupal eye leads to the formation of R7y and R7p cells. Loss of a single copy of *ss* has been shown to decrease the proportion of R7 cells that express Rh4 from 65 to 56 % (Johnston and Desplan 2014), consistent with the idea that *ss* is a rate-limiting or dosage-sensitive step in the formation of R7y cells. Cell to cell variation in gene expression may arise as the result of promoter architecture (Jones *et al.* 2014), transcriptional or translational bursting (Mcadams and Arkin 1997; Singh and Soltani 2013), cell-cell interactions (Axelrod 2010), epigenetic differences between cells (Kar *et al.* 2017) and other processes. To identify additional regulators of this process, we conducted a genetic screen to identify genetic mutations that altered the proportion of R7y and R7p cells. Here we show that the mutations in many additional genes may regulate the differentiation of R7y and R7p photoreceptor cells.

## Methods & Materials

### Drosophila stocks and genetics

All stocks were maintained in humidified incubators on standard cornmeal/molasses/agar media at 25°C. Stocks used in the experiments described here were obtained from the Bloomington Drosophila Stock Center and included *w^1118^* (FBst0003605) and *w^1118^; P{w+mW.hs=sevEP-GAL4.B}7* (FBst0005793) (*w; sevEP-GAL4*). The *sevEP-GAL4* strain was selected because it is expressed at a higher level and is more cell type specific than *P{GAL4-Hsp70.sev}* (FBtp0000379), which is driven by the Hsp70 promoter and is thought to have *sevenless* independent activity (Bailey 1999; Therrien *et al.* 1999). The 480 *P{EP}* (FBtp0001317) (Rorth 1996) transposon insertion strains used in the screen are listed in **Supplementary Table S1**.

### Screen Design

The goal of this screen was to identify genes that alter the percentage of Rh3 expressing R7 cells (Rh3%) in heterozygous mutant strains or in ectopic expression mutant strains. Each P{EP} strain was crossed to *w^1118^* flies for the heterozygous inactivation (HI) arm of the screen, and to *w; sevEP-GAL4* flies for the ectopic expression (EE) arm of the screen. For each cross 8-10 female F1 progeny were used to determine the number of R7 photoreceptor cells expressing Rh3 and Rh4 (Rh3%).

### Description of Phenotypes Scored

Immunofluorescence of R7 photoreceptor cells was performed on dissociated ommatidia, prepared as described (Hardie *et al.* 1991; Ranganathan *et al.* 1991). Rh3 and Rh4 were detected by immunohistochemistry using mouse monoclonal antibodies as previously described (Chou *et al.* 1996; Chou *et al.* 1999). Antibodies were diluted in antibody dilution buffer (3% normal goat serum, 0.03% triton X-100, and 0.1% bovine serum albumin in PBS). Dilutions used for indirect immunofluorescence were as follows: 1:20 anti-Rh3 mouse monoclonal (clone 2B1, IgG1); 1:10 anti-Rh4 mouse monoclonal (clone 11E6, IgG1). For direct immunofluorescence, directly conjugated anti-Rh3-Alexa Fluor® 488 was used at 1:50 (Earl and Britt 2006). Secondary antibodies and other immunological reagents were obtained from Jackson ImmunoResearch Laboratories, Inc. and Molecular Probes. Images were collected using an Axioskop2 Plus / AxioCamHRC microscope (Carl Zeiss, Inc.; Thornwood, NY).

### Statistical Analyses

Dissociated ommatidia were examined by fluorescence microscopy and the number of R7 cells expressing Rh3 or Rh4 were counted in each sample. To account for varying number of R7 cells examined for each cross, logistic regression with Rh3 expression (yes/no) as the outcome at the individual R7 cell-level was used to examine the association between a specific mutant in the HI screen and the probability of a cell expressing Rh3. An odds ratio (OR) was constructed to compare the odds of a cell expressing Rh3 in a specific mutant to the odds of a cell expressing Rh3 in all other mutants from the same chromosome. We adjusted for multiple testing by applying a False Discovery Rate (FDR) correction across all comparisons (all mutant strains from all chromosomes) (Benjamini and Hochberg 1995). An FDR threshold of 0.01 was used to identify mutant strains associated with the differences in the probability of an R7 cell expressing Rh3. The same model was used to determine if the probability of Rh3 expression for a specific mutant strain is significantly altered in the EE screen. Effect size is reported as a standardized percent Rh3, which is the difference between the mutant strain specific percent Rh3 and the median percent Rh3 for all mutant strains on that chromosome. Cells with inconclusive calls were not considered in the analysis.

We also examined whether the *difference* in the probability of Rh3 expression between the HI screen and the EE screen for a specific mutant strain was significantly lower/higher than other mutant strains on the same chromosome. A logistic regression model for each mutant strain was estimated using an indicator for the specific mutant strain being tested (yes/no), type of mutation (HI or EE) and the interaction effect between these two factors. The interaction effect from this model was used to determine if the effect of a particular mutation differed between the types of mutations (HI or EE). An FDR correction was used for multiple testing correction and a threshold of 0.10 was used to identify candidate genes.

#### Network Analysis

esyN (version 2.0) (Bean *et al.* 2014) was used to construct networks involved in R7 cell fate by extracting known genetic and physical interactions among candidate genes from the mutant screens and genes known to be involved in R7 cell fate specification (based on FlyBase, version FB2018_04), such as *tgo, ss and klu*. Networks were restricted to include only connections that involved at least one candidate gene.

#### Validating Candidate Genes

To validate selected candidates, we tested *P{EP}* strains as homozygotes as described above. A standardized Rh3% was calculated for the HO data and compared to the median Rh3% for each chromosome from the HI screen.

#### Research Materials and Data Availability

*Drosophila* stocks used in the experiments described here are available from the Bloomington Drosophila Stock Center. Antibodies against Rh3 and Rh4 are available on request. Data obtained in the described experiments can be found in **Supplementary Table S2** which is available at FigShare.

## Results

The genetic screen described here was developed to identify new genes involved in the cell fate decision that regulates *Rh3* versus *Rh4* expression in R7p and R7y photoreceptors cells, respectively. Three genes, *ss, tgo* and *klu* are required for this process (Wernet *et al.* 2006; Thanawala *et al.* 2013; Johnston and Desplan 2014; Anderson *et al.* 2017) and we have shown that the cell fate decision between R7p and R7y also depends upon the *Epidermal growth factor receptor* (*Egfr*) (Birkholz *et al.* 2009a). We used the *P{EP}* transposable element (Rorth 1996) in a three-part screen, taking advantage of its utility in single P-element mutagenesis and as a modular GAL4 system for over- or mis-expression (**Figure 1A-B**).

**Figure 1:**
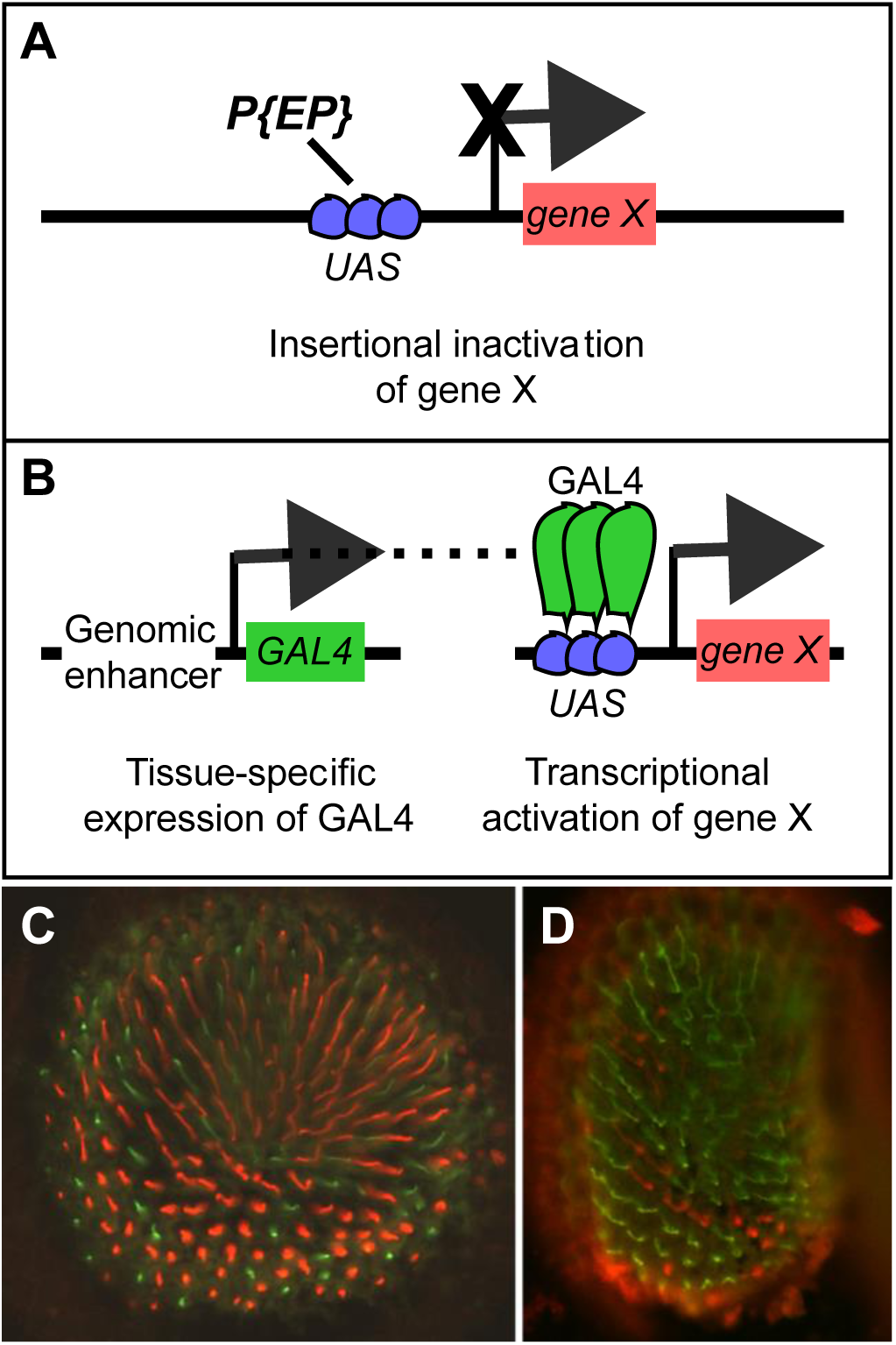
Overview of the *P{EP}* transposon insertion GAL4/UAS system and its use in this screen. **A)** The *P{EP}* transposon contains a series of yeast Upstream Activating Sequence sites (UAS, blue circles) that are the target of GAL4 binding. Upon integration into the genome in proximity to gene X, the *P{EP}* insertion may inactivate transcription of gene X or disrupt gene function by insertion elsewhere. Heterozygotes carrying such insertions were tested in the Heterozgous Insertion (HI) arm of the screen. **B)** The *sevEP-GAL4* strain (left) was crossed to the *P{EP}* insertion strain (right) to produce GAL4 in R7 photoreceptors and other cells posterior to the morphogenetic furrow, leading to transcription of gene X in a *sevEP* pattern. Animals carrying both the *sevEP-GAL4* and *P{EP}* elements were tested in the Ectopic Expression (EE) arm of the screen. **C)** The white-eyed, but otherwise wild-type *cn bw* retina with R7 cells stained for Rh3 (green) or Rh4 (red). This eye section has 31% Rh3 (*n*=232). **D)** Flies homozygous for the *slit^EP937^* insertion (identified in our screen) have 60.36% Rh3 (*n*=618). Panel A modified from (Rorth *et al.* 1998).

Our approach is based on the observation that the Rh3% of R7 photoreceptors are affected by the gene dosage of *ss* (Johnston and Desplan 2014), and the prediction that heterozygous single transposon insertions could be used to identify additional dosage sensitive loci that altered the normal 35% versus 65% percentages of Rh3 and Rh4 expressing R7 cells (**Figure 1C**). For this “heterozygous inactivation” (HI) arm of the screen, each *P{EP}* strain was crossed to *w^1118^* controls. We also hypothesized that increasing the expression of dosage sensitive loci would also alter Rh3%. For this “ectopic expression” (EE) arm of the screen, each *P{EP}* strain was crossed to *w; sevEP-GAL4.* This GAL4 strain is expressed in the pattern of *sevenless* protein and is produced dynamically in the 3^rd^ instar larval eye-antenna imaginal disc, posterior to the morphogenetic furrow in photoreceptor cells R4, R4, the mystery cells, R1, R6, R7 and the cone cells (Tomlinson *et al.* 1987; ST Pierre *et al.* 2002). An average of over 700 ommatidia were counted for each cross (range: 179-2312). The median of Rh3% for each chromosome of the HI arm of the screen are 44.0%, 38.0% and 39.1% for chromosomes X, 2 and 3 respectively, **Figure 2A**. While the median of Rh3% for each chromosome of the EE arm of the screen are 49.5%, 44.0% and 45.3% for chromosomes X, 2 and 3 respectively, **Figure 2B**. The third “difference in environments” (Diff) arm of the screen was incorporated to reveal instances where EE effects were masked by the HI, or where the observed increase / decrease of EE was due to the *P{EP}* transposon insertion itself (HI) (i.e. a comparison of the EE and HI arms of the screen). The value of Diff was calculated as Rh3% Diff = ((Rh3% in EE) – (Rh3% in HI) for the same mutant strain). The median of Rh3% Diff for each chromosome of the screen are 5.0%, 5.9% and 5.9% for chromosomes X, 2 and 3 respectively, **Figure 2C**. The medians for Rh3% for different chromosomes in all three screens (HI, EE and Diff) likely vary because of the genetics and varying sources of the chromosomes used to generate the hopped *P{EP}* strains. The Rh3% of Diff is non-zero and we believe that this results from an effect of the strains carrying a copy of only the *P{EP}* transposon in the HI screen versus carrying a copy of both the *P{EP}* and *sevEP- GAL4* transposons in the EE screen. Performing a non-parametric Kruskal-Wallis test for differences between group medians, there are significant differences across chromosomes in the median proportion of cells expressing Rh3 in all 3 screens: HI, EE and Diff screens (*p*-values 1.9 × 10^-52^, 6.3 × 10^-41^ and 6.7 × 10^-4^, respectively).

**Figure 2.**
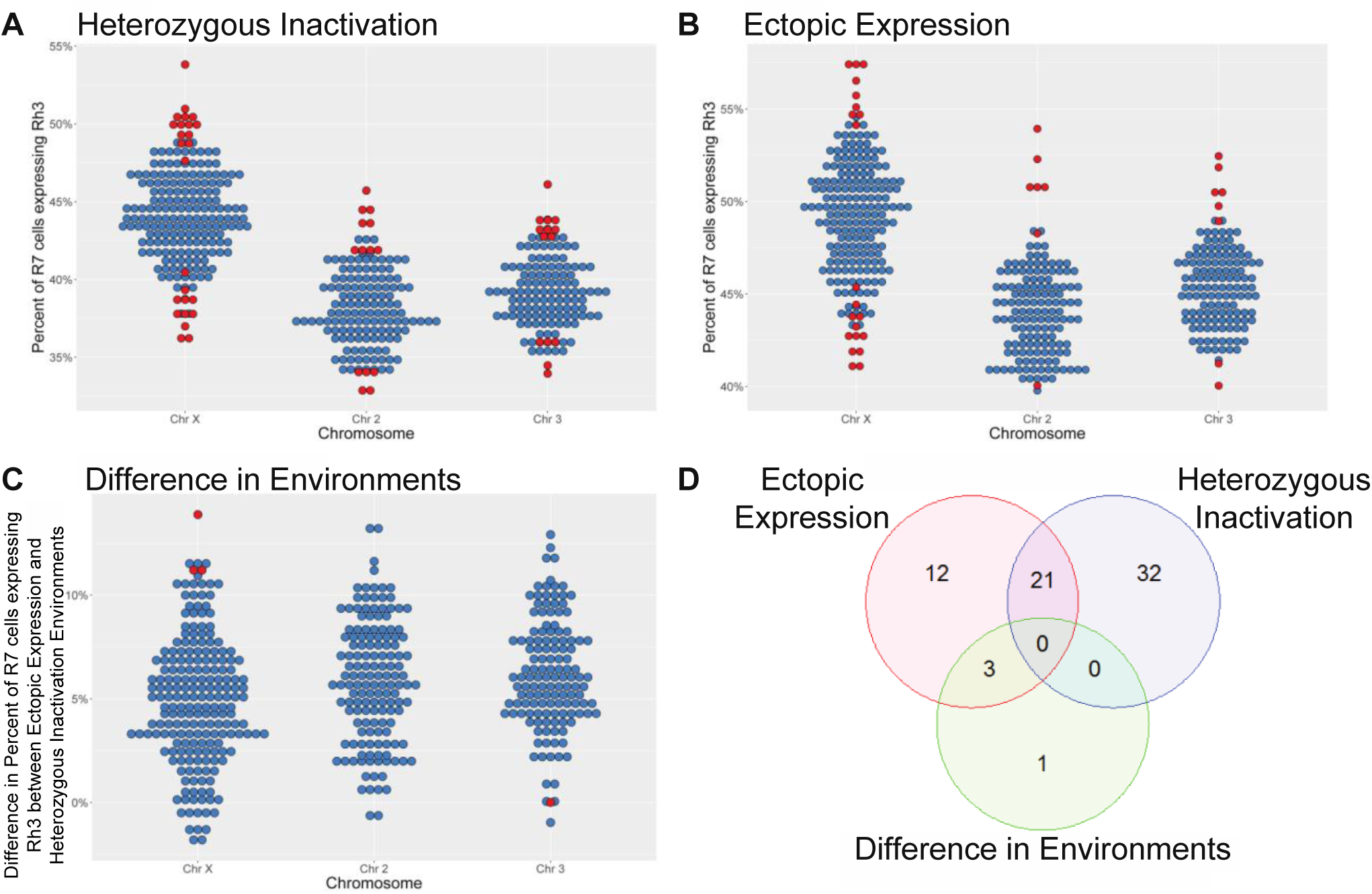
Percent of R7 cells expressing Rh3 when a single gene is altered. **A)** The panel shows the Rh3% measured for animals from the Heterozygous Insertion (HI) arm of the screen, which carry a single copy of the *P{EP}* transposon at a specific genomic location. The panel is organized by the chromosome of the insertion. Individual strains were compared to the median Rh3% for strains on that chromosome. Dots indicate individual insertion strains. Red dots correspond to measurements where the Rh3% was significantly lower / higher with a False Discovery Rate (FDR) < 0.10. Blue dots correspond to all other insertions. **B)** The panel shows the Ectopic Expression (EE) arm of the screen in which individual *P{EP}* strains were crossed to *sevEP-GAL4*. The data is presented as in Panel A. **C)** The panel shows the Difference in Environments (Diff) arm of the screen. The y-axis represents the difference in the Rh3% between the two screening environments (Diff = EE - HI) and was designed to reveal EE effects that were masked by the HI, or instances where the observed increase / decrease of EE was due to the underlying *P{EP}* insertion itself (HI). The data is presented as in Panel A. **D)** The panel shows the overlap of candidate genes among the three different arms of the screen. Each circle represents a different test. Twenty one candidates are shared between HI and EE and three candidates are shared between EE and Diff. The remaining candidates were identified in a single arm of the screen.

Based on the genomic site of the *P{EP}* insertion, we identified 53 candidate genes in the HI screen (**Figure 2A**; **Supplemental Table S2**), 36 in the EE screen (**Figure 2B**; **Supplemental Table S2**) and 4 from the Diff screen (**Figure 2C**; **Supplementary Table S2**) having an FDR <0.10. Some genes appeared in multiple screens resulting in 69 unique candidate genes in total (**Figure 2D**). Of the 53 candidate genes from the HI screen, 32 (60.4%) showed an increase in Rh3% compared to other HI mutants on the same chromosome. Likewise, of the 36 candidate genes from the EE screen, 21 (58.3%) exhibited an increase in Rh3% compared to other EE mutants from the same chromosome. Of the 4 mutants identified in the Diff arm of the screen, 3 (75%) indicated a difference in Rh3% levels between EE and HI that were greater than the Diff median for that chromosome.

For a more qualitative examination, we prioritized 10 of the 69 candidate genes (**Table 1**). These 10 include the 4 candidate genes from the Diff arm of the screen and the top 3 based on *p*-value from the HI and EE arms of the screen. In 5 of the 10 candidate genes, the HI mutant did not result in a significant change in Rh3%, but the EE screen did. In 4 of these 5, (*IGF-II mRNA-binding protein* (*Imp*), *grauzon* (*grau*), *Tyrosylprotein sulfotransferase* (*Tpst*), and *Furin 2* (*Fur2*)), Rh3% increased with EE. EE of the other gene, *Small ribonucleoprotein particle protein SmD2* (*SmD2*), led to a decrease in Rh3%. In 5 of the remaining mutant strains, two (*Tao* and *Phospolipase A2 group III (Glllspla2*)) had a significant decrease in Rh3% in the HI mutants, but the magnitude of the decrease did not change significantly with EE. Both *Actin-related protein 3* (*Arp3*) and *Rho GTPase activating protein at 18B* (*RhoGAP18B*) had a significant increase in Rh3% in their respective HI mutants, but for both mutations, that increase was dampened with EE. For both genes, the EE and Diff screens were suggestive (unadjusted p-value between 0.03 and 0.11) of a difference, but none of the four comparisons reached statistical significance after multiple testing correction. For the final gene in **Table 1**, *G protein β-subunit 13F* (*Gβ13F*), HI causes a slight up- regulation in Rh3%, while EE caused a slight down-regulation in Rh3%. Both comparisons were nominally significant but failed to pass multiple testing correction, but the Diff effect was statistically significant.

As a first step in defining the relationship between the 69 candidate genes identified in the screen and the well characterized mediators of R7 photoreceptor cell differentiation and regulation of Rh3 vs. Rh4 expression, *ss, tgo* and *klu*, we examined the known genetic and physical interactions between these genes. Of the 69 candidate genes, 66 were included in the esyN database. 28 had no known interaction in the database that was one link away from another candidate (i.e. two candidate genes were linked to the same gene), and 38 did include interactions with either another candidate(s) or an interaction with another gene 1 link away from another candidate(s). There were 85 genes that were not part of our candidate list that connected two or more candidate genes (**Figure 3**). Most candidate genes are in a single large network where the genes *expanded* (*ex*) and *pebbles* (*peb*) were the most highly connected (i.e., hub genes), while there are 2 candidate genes in a separate network. This much smaller network consisted of candidate genes *CG15514* and *jim lovell* (*lov*) connected by *MOB kinase activator 4* (*Mob4*). Only a single link is shown between a pair of genes even if there were multiple types of connections cited (e.g. a suppressing genetic interaction and a physical interaction). We identified 85 connecting genes which support connections between candidate genes that were either not evaluated in the mutant screen or were identified as statistically significant in the mutant screen. Of these 85 genes, *transcriptional Adaptor 2b* (*Ada2b*), and *scalloped* (*sd*), were included in the mutant strains screened (insertions *P{EP}EP3412* and *P{EP}sd^EP1088^*, respectively) but did not reach statistical significance. Eight candidate genes (*ex, peb, fat facets (faf), Sin3A, Imp, slit (sli), Tao* and *highwire* (*hiw)*) have connections to 10 or more other genes. **Supplementary Table S3** lists the genes in the network and the total number of connections for each gene.

**Figure 3.**
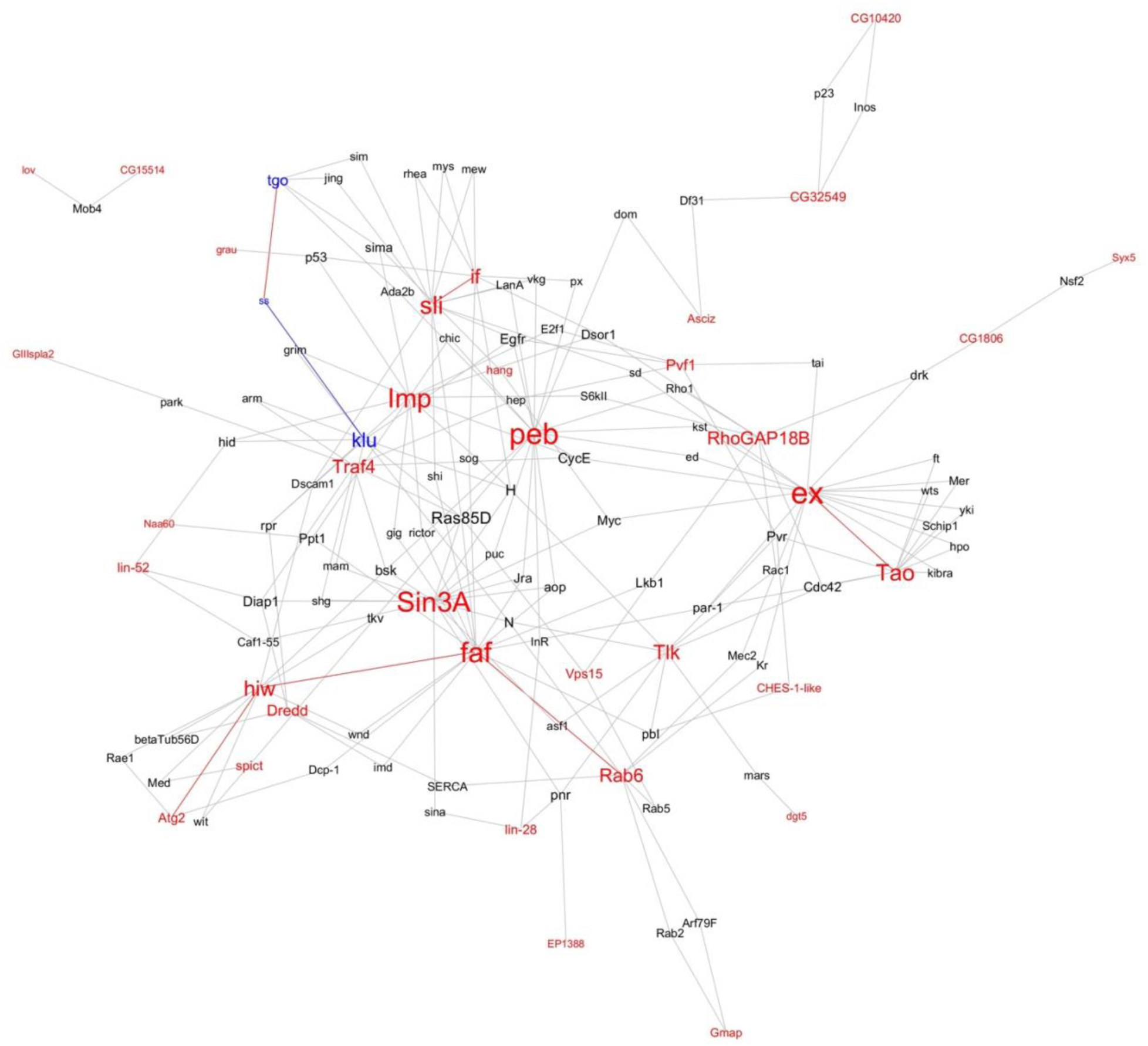
Interaction network associated with R7 cell fate differentiation and the expression of Rh3 and Rh4. Candidate genes identified in the mutant screen that alter the percent of R7 cells expressing Rh3 are shown in red. *spineless (ss), tango (tgo)* and *klumpfuss (klu)*, which are known to regulate Rh3%, are shown in blue. Genes with a known genetic or physical interaction with these genes are shown in black. Direct links between candidate genes are shown in red and manually added links between genes are shown in blue. The font size of gene name is based on the number of other genes it is connected to.

**Table 1:**
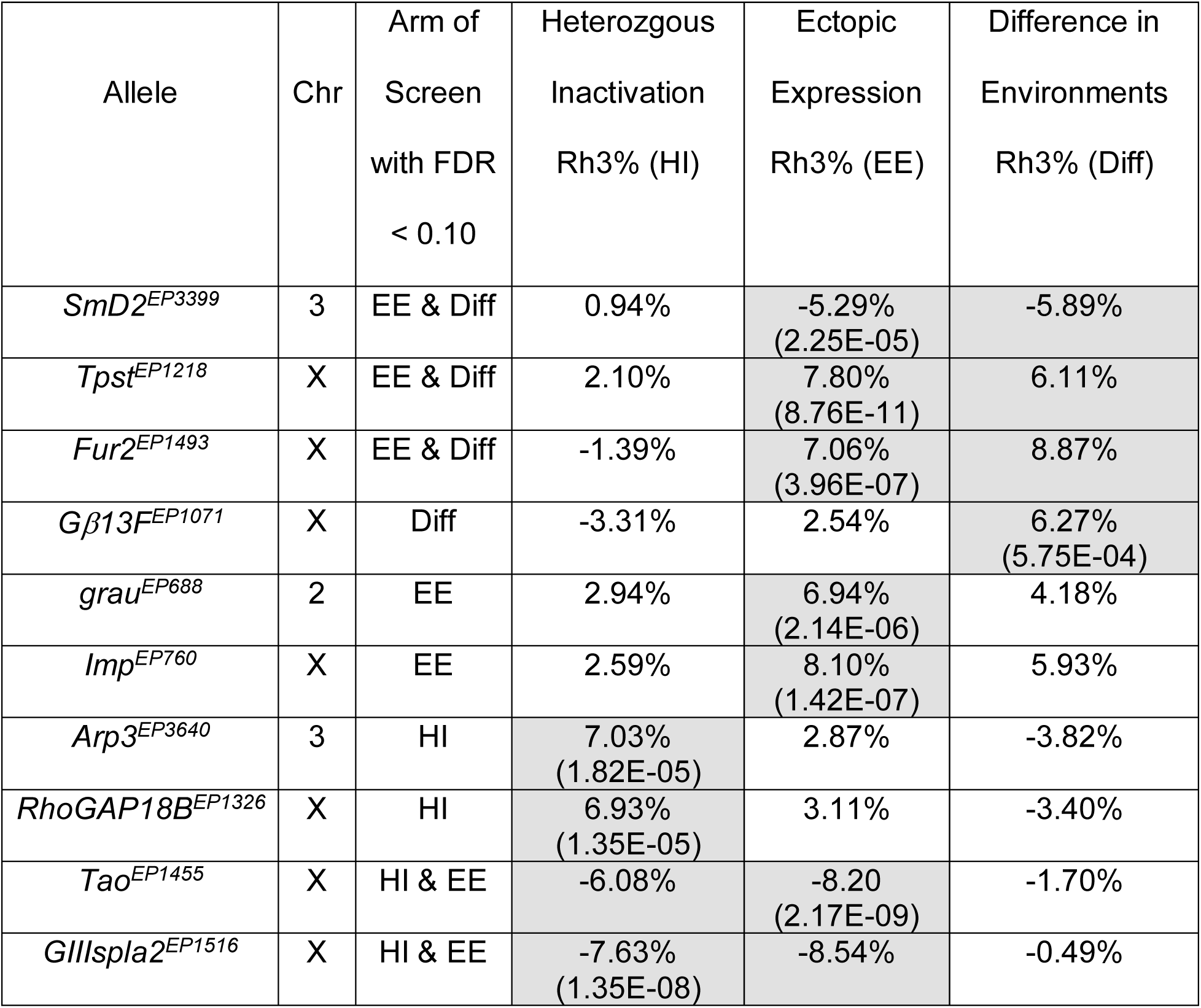
Selected Mutants with Significant Differences in R7 cells expressing Rh3. The Table shows the top candidate alleles identified in the screen, the Chromosome (Chr) containing the *P{EP}* insertion and the results of the three arms of the screen for these alleles. Abbreviations: False Discovery Rate (FDR), Standardized Percent of Rh3 expressing R7 cells (Rh3%), Heterozygous Inactivation (HI), Ectopic Expression (EE), Difference in Environments (Diff) = EE Rh3% - HI Rh3%. The table includes all 4 candidates from the Diff screen, along with the top 3 candidates from the HI and EE screens. Standardized Rh3% refers to the difference between the mutant specific Rh3% and the median Rh3% across all mutants on the same chromosome for that arm of the screen. Shaded cells indicate measurements that meet an FDR < 0.10 and the *p*-value of the highest statistical significance for that mutant strain is also shown.

As an additional step in validating the effects of mutations on Rh3%, we examined a subset of insertion strains as homozygotes. The rationale for the heterozygous insertion arm of the screen was based upon the assumption that reduction in the level of a rate-limiting regulator of R7 photoreceptor differentiation (even in heterozygotes) would demonstrate a change in Rh3%. If this was true, then our expectation is that animals homozygous for the insertion would have a further reduction in that critical regulator and would demonstrate a larger change in Rh3%. **Figure 4** shows that for the viable homozygotes tested, the change in Rh3% is more extreme than either the HI or EE state in most cases, as shown for the dramatic increase of Rh3% in *slit^EP937^* mutants (**Figure 1D** and **Figure 4**).

**Figure 4.**
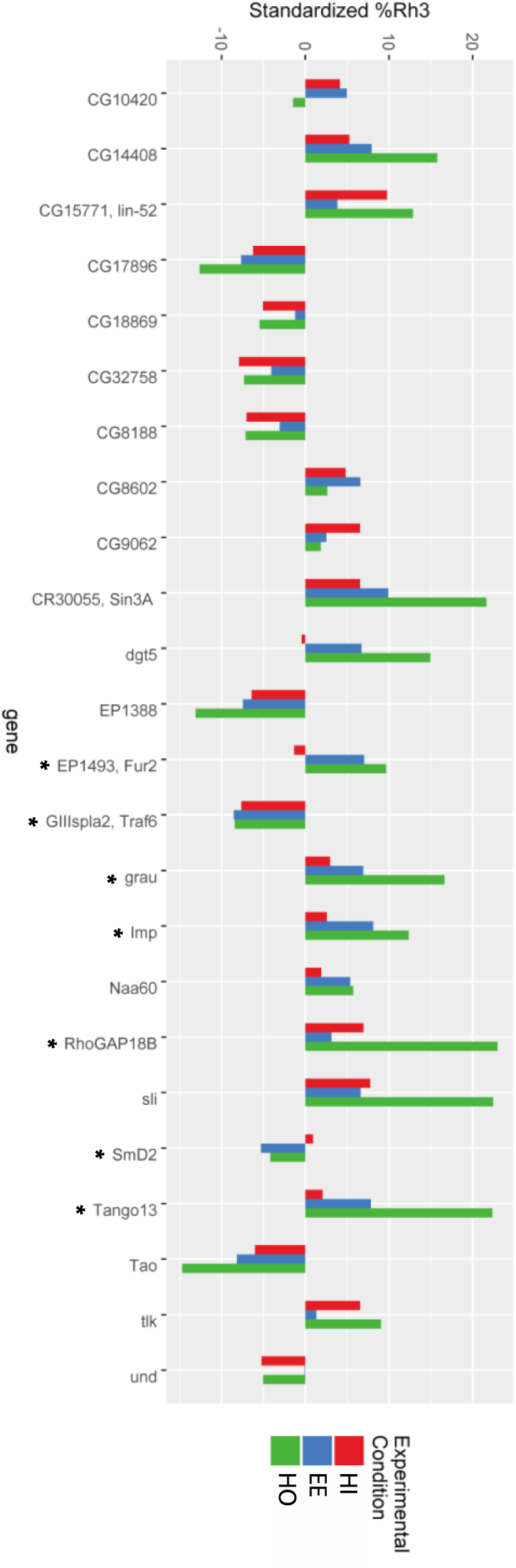
Homozygous validation of selected candidates from mutant screen. The bar graph shows the standardized Rh3% for animals in the Heterozygous Insertion (HI, red bars) and Ectopic Expression (EE, blue bars) arms of the screen. Rh3% for animals carrying Homozygous Insertions (HO) are shown in green. HO were examined for 28 selected candidates. Four of these mutants (*Arp3, Atg2, faf*, and *lov*) were homozygous lethal and are not included in the figure. A standardized Rh3% was calculated for the HO data using the Rh3% median for each chromosome from the HI screen. The asterisk (*) indicates mutants included in **Table 1**.

## Discussion

The genetic screens described here were designed to identify transposon insertion strains that showed alterations in the stochastic differentiation of R7 photoreceptor cells expressing Rh3 or Rh4. Potential alterations were defined as significant Rh3% change as transposon insertion heterozygotes, upon ectopic expression, or as a difference between the two measures. The screens identified 69 unique candidate genes. Many of the genes identified in the screen have previously identified protein-protein and genetic interaction between themselves and the known regulators of R7 cell differentiation, *klu*, *tgo* and *ss*. Preliminary validation of candidates by examination of homozygous insertions showed substantially increased effects on Rh3%.

The candidate genes identified encode proteins comprising a broad range of biological functions. Of the selected genes highlighted in **Table 1**, *SmD2* encodes an RNA binding protein that is involved in pre-mRNA splicing (Mount and Salz 2000). *Tpst* shares homology with tyrosyl-protein sulfotransferases (Gaudet *et al.* 2011), localizes to the Golgi apparatus and is involved in protein secretion (Bard *et al.* 2006). *Fur2* encodes a proprotein convertase that mediates ligand activation through protein cleavage (Kunnapuu *et al.* 2009). *Gβ13F* encodes a G-protein β-subunit that mediates G-protein coupled receptor signaling and modulates *hedgehog* signaling (Li *et al.* 2018). *grau* encodes a zinc-finger transcription factor (Chen *et al.* 2000). *Imp* encodes an mRNA binding protein that promotes and regulates transcript targeting, and plays a role in axonal remodeling, synaptogenesis and oogenesis (Boylan *et al.* 2008; Medioni *et al.* 2014). *Arp3* encodes an actin related protein that is required for myoblast fusion and axonal arborization and synapse formation (Richardson *et al.* 2007; Koch *et al.* 2014). *RhoGAP18B* encodes a GTPase activating protein that plays a role in the behavioral response to ethanol and neuromuscular junction formation (Laviolette *et al.* 2005; Rothenfluh *et al.* 2006). *Tao* encodes a mitogen-activated protein kinase kinase kinase and acts together with *hippo* (*hpo)* to activate *warts (wts)-*mediated repression of *yorkie (yki)* (Poon *et al.* 2011). *GIIIspla2* encodes a phospholipase A2 that interacts genetically with the E3 ubiquitin ligase *parkin* (Cha *et al.* 2005).

The major findings of the current work are: 1) the results provide strong support for the hypothesis that Rh3% is sensitive to dosage effects through loss of function or ectopic expression (gain of function) as might be expected for a stochastic, cell-autonomous biological process, 2) numerous genes in addition to *klu, tgo* and *ss*, are likely to play a role in regulating this process, 3) the identification of cell-cell signaling and cell surface molecules suggests that R7 cell subtype specification may not be exclusively cell-autonomous, but likely involves inputs from other cell types or tissues. Interestingly, the interaction network defined by the candidate genes contains several genes previously shown to influence specification of Rh3 or Rh4 expression in R7 photoreceptor cells (Rh3%) as well as the inductive signal that is thought to coordinate the expression of opsin genes in adjacent R7 and R8 photoreceptor cells within individual ommatidia. These include (*Egfr*) (Birkholz *et al.* 2009) that is required in both processes, as well as *Merlin* (*Mer*), *wts*, *hpo*, *kibra yki* (Jukam and Desplan 2011; Jukam *et al.* 2013) and *thickveins* (*tkv*) (Wells *et al.* 2017) that coordinate opsin gene expression in R7 and R8 cells. Furthermore, one of the candidate genes identified in the screen, *Fur2*, has also been shown to regulate the induction of Rh5 expression in R8 cells (Wells *et al.* 2017). These results suggest numerous mechanisms through which processes in the R7 cell responsible for R7y versus R7p cell fate may couple to inductive signaling that specifies paired R8y and R8p cell fates.

The initial analyses here are insufficient to determine the precise mechanism of action of the identified candidate genes. They could potentially function cell autonomously within the R7 photoreceptor cell genetically upstream (and epistatic to) or down stream of *ss*. Alternatively, the candidate genes may also play roles outside of the R7 cell and function independently of *ss*. Finally, the genes identified in this study may play an instructive versus a permissive role in R7 cell differentiation. Additional experiments will be required to resolve these mechanistic questions.

## Supplementary Tables

**S1. Table of *P{EP}* mutants used in the screen.** The columns indicate the insertion name, Flybase transposon insertion code (FBti) and the Bloomington Drosophila Stock Center stock number and genotype (if available).

**S2: Table of Full Screen Results.** The columns indicate the chromosome carrying the insertion, the *P{EP}* line, the gene effected, Raw Data Counts for Rh4 and Rh3 expression in the Heterozygous Inactivation (HI) and Ectopic Expression (EE) arms of the screen, the calculated Rh3% for HI and EE and the calculated Difference between the two environments (Diff = EE – HI). Logistic Regression Testing results of HI, EE and Diff are shown that each list WaldChiSq, p-value, and FDR (False Discovery Rate). The table is sorted by increasing FDR for Diff, EE and HI. FDR < 0.05 is shaded in yellow. FDR < 0.10 is shaded in gray.

**S3: Interaction Network Details.** The table provides data on the Interaction Network described in **Figure 3**. The columns indicate the genes within the network, whether they are candidates identified in the screen, the total number of connections and the number of unique connections to that gene.

## References

Anderson, C., I. Reiss, C. Zhou, A. Cho, H. Siddiqi et al., 2017 Natural variation in stochastic photoreceptor specification and color preference in Drosophila. Elife 6.

Axelrod, J. D., 2010 Delivering the lateral inhibition punchline: it’s all about the timing. Sci Signal 3: pe38.

Bailey, A., 1999 sevenless-GAL4 transgene. Personal Communication to FlyBase. FBrf0125052

Bard, F., L. Casano, A. Mallabiabarrena, E. Wallace, K. Saito et al., 2006 Functional genomics reveals genes involved in protein secretion and Golgi organization. Nature 439: 604–607.

Bean, D. M., J. Heimbach, L. Ficorella, G. Micklem, S. G. Oliver et al., 2014 esyN: network building, sharing and publishing. PLoS One 9: e106035.

Bell, M. L., J. B. Earl and S. G. Britt, 2007 Two types of Drosophila R7 photoreceptor cells are arranged randomly: a model for stochastic cell-fate determination. J Comp Neurol 502: 75–85.

Benjamini, Y., and Y. Hochberg, 1995 Controlling the False Discovery Rate: A Practical and Powerful Approach to Multiple Testing. Journal of the Royal Statistical Society. Series B (Methodological) 57: 289–300.

Birkholz, D. A., W. H. Chou, M. M. Phistry and S. G. Britt, 2009 rhomboid mediates specification of blue- and green-sensitive R8 photoreceptor cells in Drosophila. J Neurosci 29: 2666–2675.

Boylan, K. L., S. Mische, M. Li, G. Marques, X. Morin et al., 2008 Motility screen identifies Drosophila IGF-II mRNA-binding protein--zipcode-binding protein acting in oogenesis and synaptogenesis. PLoS Genet 4: e36.

Cha, G. H., S. Kim, J. Park, E. Lee, M. Kim et al., 2005 Parkin negatively regulates JNK pathway in the dopaminergic neurons of Drosophila. Proc Natl Acad Sci U S A 102: 10345–10350.

Chen, B., E. Harms, T. Chu, G. Henrion and S. Strickland, 2000 Completion of meiosis in Drosophila oocytes requires transcriptional control by grauzone, a new zinc finger protein. Development 127: 1243–1251.

Chou, W. H., K. J. Hall, D. B. Wilson, C. L. Wideman, S. M. Townson et al., 1996 Identification of a novel Drosophila opsin reveals specific patterning of the R7 and R8 photoreceptor cells. Neuron 17: 1101–1115.

Chou, W. H., A. Huber, J. Bentrop, S. Schulz, K. Schwab et al., 1999 Patterning of the R7 and R8 photoreceptor cells of Drosophila: evidence for induced and default cell-fate specification. Development 126: 607–616.

Cook, T., F. Pichaud, R. Sonneville, D. Papatsenko and C. Desplan, 2003 Distinction between color photoreceptor cell fates is controlled by Prospero in Drosophila. Dev Cell 4: 853–864.

Earl, J. B., and S. G. Britt, 2006 Expression of Drosophila rhodopsins during photoreceptor cell differentiation: Insights into R7 and R8 cell subtype commitment. Gene Expr Patterns 6: 687–694.

Fortini, M. E., and G. M. Rubin, 1991 The optic lobe projection pattern of polarization-sensitive photoreceptor cells in Drosophila melanogaster. Cell Tissue Res 265: 185–191.

Franceschini, N., K. Kirschfeld and B. Minke, 1981 Fluorescence of photoreceptor cells observed in vivo. Science 213: 1264–1267.

Gaudet, P., M. S. Livstone, S. E. Lewis and P. D. Thomas, 2011 Phylogenetic-based propagation of functional annotations within the Gene Ontology consortium. Brief Bioinform 12: 449–462.

Hardie, R. C., D. Voss, O. Pongs and S. B. Laughlin, 1991 Novel potassium channels encoded by the Shaker locus in Drosophila photoreceptors. Neuron 6: 477–486.

Jacobs, G. H., 1981 Comparative color vision. Academic Press, New York.

Johnston, R. J., Jr., and C. Desplan, 2014 Interchromosomal communication coordinates intrinsically stochastic expression between alleles. Science 343: 661–665.

Jones, D. L., R. C. Brewster and R. Phillips, 2014 Promoter architecture dictates cell-to-cell variability in gene expression. Science 346: 1533–1536.

Jukam, D., and C. Desplan, 2011 Binary regulation of Hippo pathway by Merlin/NF2, Kibra, Lgl, and Melted specifies and maintains postmitotic neuronal fate. Dev Cell 21: 874–887.

Jukam, D., B. Xie, J. Rister, D. Terrell, M. Charlton-Perkins et al., 2013 Opposite feedbacks in the Hippo pathway for growth control and neural fate. Science 342: 1238016.

Kar, G., J. K. Kim, A. A. Kolodziejczyk, K. N. Natarajan, E. Torlai Triglia et al., 2017 Flipping between Polycomb repressed and active transcriptional states introduces noise in gene expression. Nat Commun 8: 36.

Kirschfeld, K., R. Feiler and N. Franceschini, 1978 A photostable pigment within the rhabdomeres of fly photoreceptors no. 7. J. Comp. Physiol. 125: 275–284.

Koch, N., O. Kobler, U. Thomas, B. Qualmann and M. M. Kessels, 2014 Terminal axonal arborization and synaptic bouton formation critically rely on abp1 and the arp2/3 complex. PLoS One 9: e97692.

Kunnapuu, J., I. Bjorkgren and O. Shimmi, 2009 The Drosophila DPP signal is produced by cleavage of its proprotein at evolutionary diversified furin-recognition sites. Proc Natl Acad Sci U S A 106: 8501–8506.

Laviolette, M. J., P. Nunes, J. B. Peyre, T. Aigaki and B. A. Stewart, 2005 A genetic screen for suppressors of Drosophila NSF2 neuromuscular junction overgrowth. Genetics 170: 779–792.

Li, S., Y. S. Cho, B. Wang, S. Li and J. Jiang, 2018 Regulation of Smoothened ubiquitylation and cell surface expression through a Cul4-DDB1-Gbeta E3 ubiquitin ligase complex. J Cell Sci 131.

McAdams, H. H., and A. Arkin, 1997 Stochastic mechanisms in gene expression. Proc Natl Acad Sci U S A 94: 814–819.

Medioni, C., M. Ramialison, A. Ephrussi and F. Besse, 2014 Imp promotes axonal remodeling by regulating profilin mRNA during brain development. Curr Biol 24: 793–800.

Mikeladze-Dvali, T., M. F. Wernet, D. Pistillo, E. O. Mazzoni, A. A. Teleman et al., 2005 The Growth Regulators warts/lats and melted Interact in a Bistable Loop to Specify Opposite Fates in Drosophila R8 Photoreceptors. Cell 122: 775–787.

Mount, S. M., and H. K. Salz, 2000 Pre-messenger RNA processing factors in the Drosophila genome. J Cell Biol 150: F37–44.

Nathans, J., D. Thomas and D. S. Hogness, 1986 Molecular genetics of human color vision: the genes encoding blue, green, and red pigments. Science 232: 193–202.

Papatsenko, D., G. Sheng and C. Desplan, 1997 A new rhodopsin in R8 photoreceptors of Drosophila: evidence for coordinate expression with Rh3 in R7 cells. Development 124: 1665–1673.

Poon, C. L., J. I. Lin, X. Zhang and K. F. Harvey, 2011 The sterile 20-like kinase Tao-1 controls tissue growth by regulating the Salvador-Warts-Hippo pathway. Dev Cell 21: 896–906.

Quinn, W. G., W. A. Harris and S. Benzer, 1974 Conditioned behavior in Drosophila melanogaster. Proc Natl Acad Sci U S A 71: 708–712.

Ranganathan, R., W. A. Harris and C. S. Zuker, 1991 The molecular genetics of invertebrate phototransduction. Trends in Neurosciences 14: 486–493.

Richardson, B. E., K. Beckett, S. J. Nowak and M. K. Baylies, 2007 SCAR/WAVE and Arp2/3 are crucial for cytoskeletal remodeling at the site of myoblast fusion. Development 134: 4357–4367.

Rorth, P., 1996 A modular misexpression screen in Drosophila detecting tissue-specific phenotypes. Proc Natl Acad Sci U S A 93: 12418–12422.

Rorth, P., K. Szabo, A. Bailey, T. Laverty, J. Rehm et al., 1998 Systematic gain-of-function genetics in Drosophila. Development 125: 1049–1057.

Rothenfluh, A., R. J. Threlkeld, R. J. Bainton, L. T. Tsai, A. W. Lasek et al., 2006 Distinct behavioral responses to ethanol are regulated by alternate RhoGAP18B isoforms. Cell 127: 199–211.

Singh, A., and M. Soltani, 2013 Quantifying intrinsic and extrinsic variability in stochastic gene expression models. PLoS One 8: e84301.

Spatz, H. C., A. Emanns and H. Reichert, 1974 Associative learning of Drosophila melanogaster. Nature 248: 359–361.

St Pierre, S. E., M. I. Galindo, J. P. Couso and S. Thor, 2002 Control of Drosophila imaginal disc development by rotund and roughened eye: differentially expressed transcripts of the same gene encoding functionally distinct zinc finger proteins. Development 129: 1273–1281.

Tang, S., and A. Guo, 2001 Choice behavior of Drosophila facing contradictory visual cues. Science 294: 1543–1547.

Thanawala, S. U., J. Rister, G. W. Goldberg, A. Zuskov, E. C. Olesnicky et al., 2013 Regional modulation of a stochastically expressed factor determines photoreceptor subtypes in the Drosophila retina. Dev Cell 25: 93–105.

Therrien, M., A. M. Wong, E. Kwan and G. M. Rubin, 1999 Functional analysis of CNK in RAS signaling. Proc Natl Acad Sci U S A 96: 13259–13263.

Tomlinson, A., D. D. Bowtell, E. Hafen and G. M. Rubin, 1987 Localization of the sevenless protein, a putative receptor for positional information, in the eye imaginal disc of Drosophila. Cell 51: 143–150.

Viets, K., K. C. Eldred and R. J. Johnston, Jr., 2016 Mechanisms of Photoreceptor Patterning in Vertebrates and Invertebrates. Trends Genet 32: 638–659.

Wells, B. S., D. Pistillo, E. Barnhart and C. Desplan, 2017 Parallel activin and BMP signaling coordinates R7/R8 photoreceptor subtype pairing in the stochastic Drosophila retina. Elife 6.

Wernet, M. F., T. Labhart, F. Baumann, E. O. Mazzoni, F. Pichaud et al., 2003 Homothorax switches function of Drosophila photoreceptors from color to polarized light sensors. Cell 115: 267–279.

Wernet, M. F., E. O. Mazzoni, A. Celik, D. M. Duncan, I. Duncan et al., 2006 Stochastic spineless expression creates the retinal mosaic for colour vision. Nature 440: 174–180.

Wikler, K. C., and P. Rakic, 1990 Distribution of photoreceptor subtypes in the retina of diurnal and nocturnal primates. J Neurosci 10: 3390–3401.

Yamaguchi, S., C. Desplan and M. Heisenberg, 2010 Contribution of photoreceptor subtypes to spectral wavelength preference in Drosophila. Proc Natl Acad Sci U S A 107: 5634–5639.

